# Trans-segmental imaging in the spinal cord of behaving mice

**DOI:** 10.1101/2021.12.23.474042

**Authors:** Pavel Shekhtmeyster, Daniela Duarte, Erin M. Carey, Alexander Ngo, Grace Gao, Jack A. Olmstead, Nicholas A. Nelson, Axel Nimmerjahn

## Abstract

Spinal cord circuits play crucial roles in transmitting and gating cutaneous somatosensory modalities, such as pain, but the underlying activity patterns within and across spinal segments in behaving mice have remained elusive. To enable such measurements, we developed a wearable widefield macroscope with a 7.9 mm^2^ field of view, subcellular lateral resolution, 2.7 mm working distance, and <10 g overall weight. We show that highly localized painful mechanical stimuli evoke widespread, coordinated astrocyte excitation across multiple spinal segments.

## Introduction

Spinal cord circuits are indispensable for sensory information processing and controlling reflexive and adaptive behaviors, such as those driven by pain and itch, as demonstrated by genetic, anatomical, electrophysiological, pharmacologic, and behavioral methods^1^. However, these approaches lack the spatiotemporal resolution, resilience against tissue movement, or cell-type specificity necessary to interrogate genetically defined cell types’ activity patterns in behaving animals. In vivo fluorescence imaging overcomes many of these barriers by permitting high-speed, stable measurement of neuronal and non-neuronal activity in the spinal cord^2^.

Sensory stimuli can recruit cellular networks distributed across multiple spinal segments^2,3^ (each ∼1-1.5 mm rostra-caudal length in mice^4^). Current fluorescence microscopes’ optical properties hamper the high-speed measurement of such distributed activity. While cellular-resolution devices typically offer small fields of view (FOVs) covering less than one vertebra, large-FOV microscopes (“macroscopes”) provide limited spatiotemporal resolution, sensitivity, and contrast, or are incompatible with use in freely moving mice^2,5,6^. Wearable widefield macroscopes with millimeter-sized FOV have recently been developed^7-9^. Still, none offer the cellular resolution, frame rates, or weights necessary for spinal cord imaging in mice. These technical limitations hamper our ability to decipher how broadly dispersed cellular networks contribute to somatosensory processing in behaving animals. Of particular interest, astrocytes are emerging as essential regulators of pain signaling in the spinal dorsal horn^10-13^.

To address these challenges, we developed a wearable macroscope with custom-compound microlenses that allows subcellular-resolution, high-speed activity measurements from genetically defined cell types across multiple spinal segments (7.9 mm^2^ field of view) in behaving animals. Using GFAP-GCaMP6f mice with calcium indicator expression in spinal astrocytes, we show that even highly localized noxious mechanical stimuli can evoke widespread, coordinated calcium activity that spans multiple spinal segments. In contrast, innocuous stimuli to the same cutaneous area led to only sparsely distributed calcium excitation, suggesting categorical differences in these activity patterns’ underlying cellular and molecular mechanisms.

## Results

### A wearable widefield macroscope with custom-compound microlenses for millimeter-scale, subcellular-resolution measurements in freely behaving mice

To meet the application needs mentioned above, the design goals for our macroscope’s optical system included an ∼3.2 × 2.5 mm FOV, high spatial resolution and contrast across the FOV (∼ 3 μm), a long working distance (> 2 mm), and broad achromatic range (450-570 nm). To provide flexibility in the type and the number of light sources that can be coupled (e.g., DPSS lasers, LEDs), a flexible PMMA fiber was chosen for excitation light delivery (**Fig. 1a-b**). Additional design parameters included an ∼0.4 numerical aperture, ∼1.4x magnification, and 36 mm track length. To ensure that mice can comfortably wear the macroscope on its back, we aimed for an overall weight of <10 g (including all optics, optomechanical components, and electronics). To minimize torque on the animal, we also opted for a folded imaging path (i.e., the short-pass dichroic beamsplitter reflects fluorescence emission and transmits excitation light), thereby lowering the integrated device’s center of gravity.

**Fig. 1.**
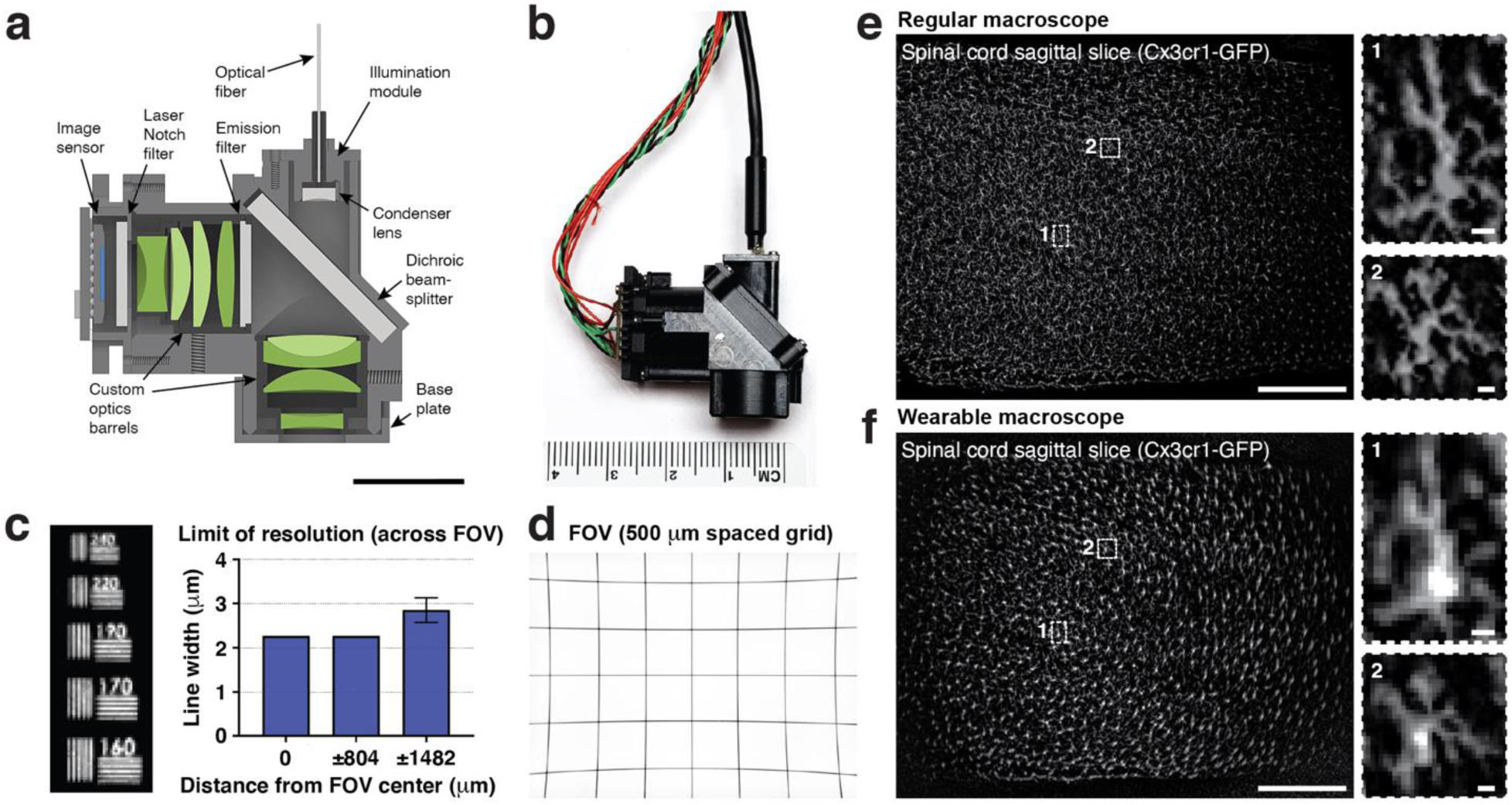
Wearable macroscopes with custom-compound-microlenses for millimeter-scale, subcellular-resolution measurements in freely behaving mice. **a**, Cross-section of the wearable macroscope and its custom compound micro-optics (green). Scale bar, 10 mm. **b**, Photo of the fabricated device. **c**, Limit of resolution across the FOV. Displayed values are averages across the horizontal and vertical line target results from comparable positions to the left and right of the FOV center. Spatial frequencies were converted to line widths. The data are presented as mean ± s.e.m. The error bars at 0 μm and ±804 μm are too small to be visible. **d**, Image of a 500-μm spacing grid target acquired with the integrated device demonstrating its ∼3.2 × 2.5 mm FOV. **e**, *Left*, fluorescence image of a 20-μm-thick sagittal spinal cord section from a Cx3cr1^+/GFP^ mouse with labeled microglia acquired with a commercial benchtop macroscope. *Right*, blow-up of the two subregions indicated on the left showing individual microglial cells. **f**, *Left*, fluorescence image of the same spinal cord section as in c acquired with the wearable macroscope. *Right*, blow-up of the subregions indicated on the left. Over 1,000 individual microglial cells are visible. Scale bars in e and f, 500 μm (left) and 10 μm (right).

Our optical design included eight custom microlenses from five different glass materials (**Suppl. Table 1**). All glasses were high-index (n=1.76 - 1.92) to increase optical power while maintaining device compactness (**Fig. 1a-b**). Zemax modeling was used to optimize the lens surface profile, radius of curvature, distance between surfaces, and glass types. We included a #0 coverslip (∼100 μm thickness) in our design to minimize optical aberrations when imaging through a live animal-implanted glass window. Additionally, we accounted for optical filters placed in the collimated space between the lenses required for fluorescence imaging. Following optical modeling, the lenses and brass spacers were fabricated and assembled into two custom-made barrel holders minimized in size and weight (**Fig. 1a-b**). Each assembled barrel weighed <2.05 g.

Next, we characterized our integrated system’s optical performance by determining its limit of resolution (LOR) across and point spread function (PSF) in the center of the FOV. For the LOR, we measured 2.27 μm in the center and halfway toward the edges of the FOV, and 2.85 μm at the FOV edges (**Fig. 1c**). The lateral and axial point spread function (PSF) of our integrated system was ∼3.3 μm and ∼17.7 μm, respectively, measured as full width half maximum (FWHM) (**Suppl. Fig. 1**). Because the ability to retrieve image information can be limited by contrast, independent of resolution, we quantified our macroscope’s modulation transfer function (MTF). Our integrated system’s MTF10 (i.e., contrast at 10%; **Methods**) was 169 lp/mm tangential (2.96 μm linewidth) and 193 lp/mm sagittal (2.59 μm linewidth) in the center of the FOV (**Suppl. Fig. 2**). Notably, our macroscope performs well even at the FOV edges (**Fig. 1f**). The ability to obtain high-contrast measurements becomes especially important in densely labeled tissue (**Figs. 1e-f, 2b, d**).

Using a precision differential actuator and regularly spaced grid target with 500 μm spacing, we found that our wearable device had a 3.215 × 2.465 mm FOV (**Fig. 1d**) and 2.727 mm working distance (WD) (from the edge of the objective barrel to the microscopy target in air). This long WD enables imaging through implantable micro-optics, such as microprisms (**Suppl. Fig. 3**). Our fully assembled device measured ∼14 × 26 × 29 mm (4.423 cm^3^ volume) and weighed 9.8 g. Remarkably, 7-9 weeks-old mice could carry the device on their back without training. Open-field measurements confirmed that general locomotor activity (average running velocity, total distance traveled, rearing activity) was largely unaffected (**Suppl. Fig. 4**).

Together, these data demonstrate that our miniature macroscope with custom-compound-microlenses simultaneously provides an exceptional field of view, spatial resolution, contrast, and working distance while retaining a small form factor and low weight suitable for spinal cord imaging in freely moving mice.

### High-speed trans-segmental imaging of sensory-evoked calcium activity in behaving GFAP-GCaMP6f mice

To date, studies investigating somatosensory processing have been limited to individual spinal segments (each ∼1-1.5 mm length in mice) or less, given previous microscopes’ technical limitations. As a result, little is known about the spatial extent and temporal evolution of single-trial activity patterns across spinal segments.

To demonstrate the feasibility of high-speed, trans-segmental measurements with our wearable macroscopes, we performed imaging in awake GFAP-GCaMP6f mice (N=4) with constitutive green-fluorescent calcium indicator GCaMP6f expression in astrocytes. Dorsal horn astrocytes can exhibit noxious stimulus-evoked activity across large tissue regions (>500 μm rostrocaudal extent, limited by FOV)^10,11^, and this activity has been suggested to serve pain modulatory functions^11-13^. However, its spatial extent and temporal dynamics across spinal segments remain unknown. Determining the spatiotemporal properties of astrocytes’ excitation is crucial to help define its functional role (e.g., trans-segmental and -hemispheric sensitization), underlying mechanisms (e.g., local or neuromodulatory signaling), and interventions to control it (e.g., through local or subpial viral vector injections)^14^. Using a rodent pincher system, we applied quantifiable and repeatable pressure stimuli to the animal’s proximal tail (duration: 1.5 s ± 0.5 s; pressure range: 50 - 1,100 g) (**Methods**) while imaging astrocytes’ calcium excitation across L2-L5 spinal segments at 25-75 μm focal depths (**Fig. 2a**). To facilitate the readout of mouse locomotor activity, we placed the mice on a spherical treadmill during imaging. To quantify sensory-evoked astrocyte responses comprehensively, we tiled the FOV with equally sized (∼10 μm × 10 μm) ROIs, excluding blood vessel regions (**Methods**).

**Fig. 2.**
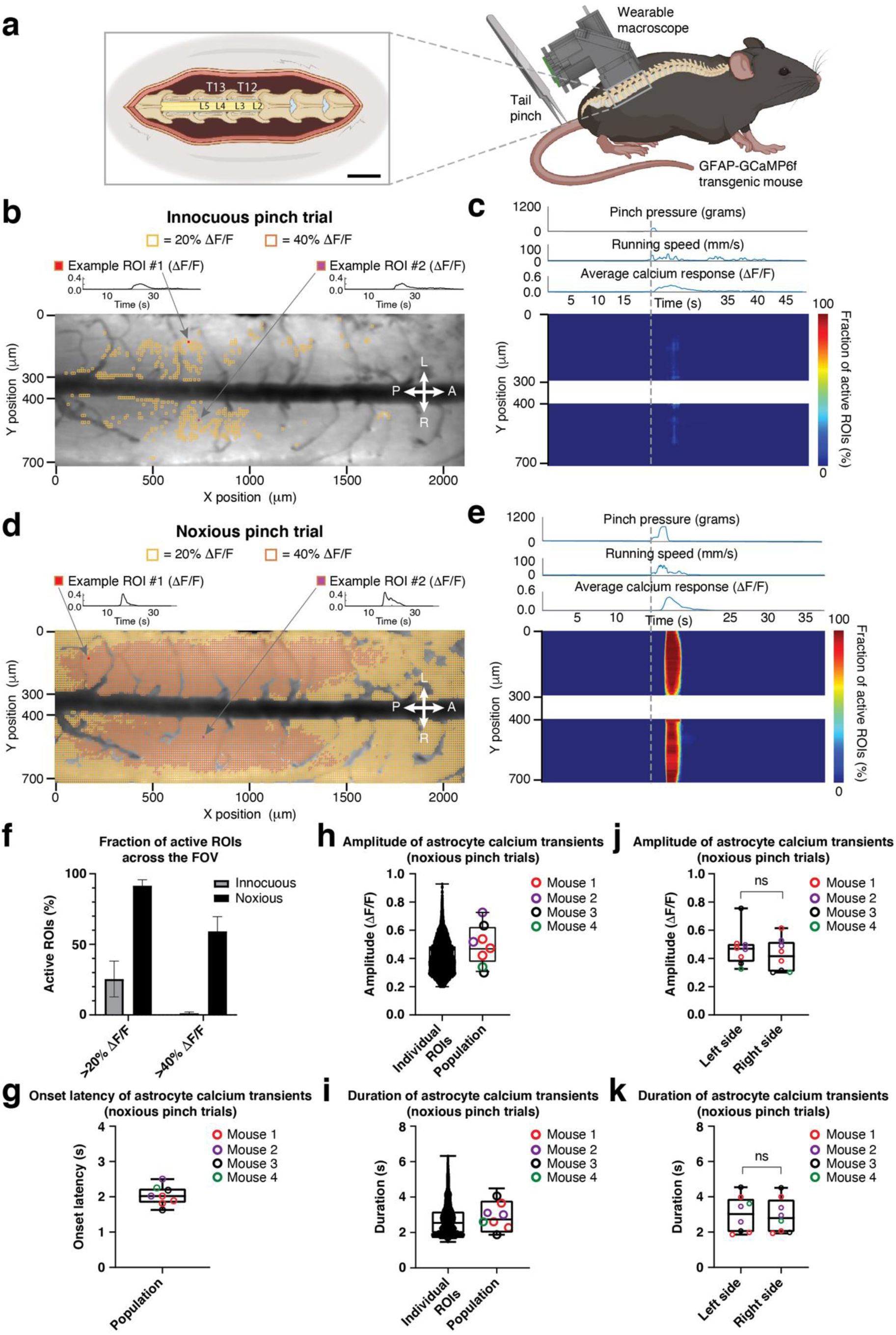
High-speed trans-segmental imaging of sensory-evoked calcium activity in behaving GFAP-GCaMP6f mice. **a**, Schematic of the experimental approach. A laminectomy was performed over the lumbar spinal cord of GFAP-GCaMP6f mice (N=4). The wearable macroscope was mounted above it. Pinch stimuli of different intensities and set duration (1.5 s ± 0.5 s) were delivered to the awake animal’s proximal tail. Running speed was recorded by placing the animal on a spherical treadmill (**Methods**). Left inset scale bar, 1,400 μm. **b**, Average intensity projection image from a time-lapse recording taken at ∼50 μm focal depth below the pia overlaid with ∼10 μm × 10 μm ROIs. Only ROIs with at least 20% (yellow) and 40% ΔF/F (orange) in response to an innocuous pinch (p < 200 g) are shown. Innocuous stimuli evoked only sparse astrocyte activity on the left (L) and right (R) sides of the spinal cord and across anterior (A) and posterior (P) lumbar spinal segments (**Suppl. Movie 1**). Two example ROI’s calcium transients are shown above the activity map. **c**, Innocuous tail pinch-evoked activity for the example recording shown in b. Each row depicts the percent of active ROIs (>20% ΔF/F) across a given mediolateral (Y) position. The corresponding pressure stimulus, locomotor activity, and average calcium transient across the FOV are shown above the activity heat map. **d**, Activity map from the same animal and focal depth as in b for a noxious pinch trial (p > 500 g). Noxious stimuli evoked widespread, bi-lateral astrocyte excitation across spinal segments (**Suppl. Movie 2**). Two example ROI’s calcium transients are shown above the activity map. **e**, Noxious tail pinch-evoked astrocyte activity across different Y positions. Pressure stimulus amplitude, locomotor activity, and average calcium signal across the FOV are shown above the activity heat map. **f**, Population data showing the percent of active ROIs with at least 20% or 40% ΔF/F for innocuous and noxious stimulus trials. **g**, Population data showing the average calcium transient onset latency for noxious pinch trials. **h-i**, Population data showing the individual ROI and average calcium response amplitude (h) and duration (i) for noxious pinch trials. **j-k**, Population data showing the average calcium transient amplitude (j) and duration (k) on the left and right sides of the spinal cord for noxious pinch trials. The data in f are from 9 innocuous and 8 noxious pinch trials in 4 mice. The data in h (left) are from 67,696 ROIs with ΔF/F >20%, 8 recordings, and 4 mice. The data in i (left) are from 62,330 ROIs with ΔF/F >40%, 8 recordings, and 4 mice. The data in g, h-i (right), and j-k are from 8 recordings and 4 mice. Paired t-tests determined *P* values, and all bar plots are presented as mean ± s.e.m. The box and violin plots mark the median and the 25th and 75th percentiles, and the whiskers cover the minimum and maximum of the data.

Innocuous tail pinch stimuli (p < 200 g) led to only sparse, low amplitude transients (most <20% ΔF/F) along the length of the >2,000 μm laminectomy (**Fig. 2b-c**; **Suppl. Movie 1**). In contrast, noxious tail pinch (p > 500 g) evoked widespread coordinated calcium transients bilaterally and across spinal segments, with many transients exceeding 40% ΔF/F (**Fig. 2d-f**; **Suppl. Movie 2**). While noxious stimulus application often evoked locomotor activity, running in the absence of tail pinch did not evoke comparable population activity (**Fig. 2c, e**; **Suppl. Movie 1**) (**Methods**), consistent with previous work^10^. Calcium transients evoked by noxious pinch had an onset latency of 2.04 ± 0.07 s (population average) (**Fig. 2g**), amplitude of 0.41 ± 0.01 ΔF/F (individual ROIs) and 0.49 ± 0.14 ΔF/F (population average) (**Fig. 2h**), and duration of 2.70 ± 0.01 s (individual ROIs) and 2.88 ± 0.97 s (population average) (**Fig. 2i**). The amplitude and duration of population transients were comparable on the left and right sides of the spinal cord (**Fig. 2j, k**) (**Methods**). Based on onset latency, the activity originated in central gray matter regions of each hemisphere (**Fig. 2e**). Together, these data demonstrate that our wearable macroscopes are capable of high-resolution, high-speed bilateral and trans-segmental measurement of sensory-evoked activity in genetically defined cells of behaving mice.

## Discussion

The technical capability of our wearable macroscopes for millimeter-scale, subcellular-resolution measurements was enabled by custom-compound microlenses (**Fig. 1a, c-d, f**). These custom optics have a theoretical lateral and axial PSF of 0.55 μm and 9.03 μm, respectively, based on Zemax calculations. Given the experimentally measured lateral and axial resolution of ∼2.3 - 3.3 μm and 17.7 μm, respectively, this suggests that our wearable macroscopes are currently limited by the image sensor’s resolution (∼2.63 μm in the object space). Therefore, future CMOS sensor upgrades are expected to improve the device’s overall resolution even further. Our device’s modular design greatly facilitates such upgrades (**Methods**).

**Fig. 2** demonstrates our wearable macroscopes’ ability to perform high-speed, high-sensitivity functional measurements from genetically defined cells across multiple spinal segments. The properties of tail pinch- and running-evoked calcium transients in GFAP-GCaMP6f mice (e.g., onset latency, duration) were consistent with previous studies^2,10^. Notably, the significantly larger FOV allowed us to reveal that even focally delivered sensory stimuli could evoke remarkably widespread, trans-segmental calcium excitation in dorsal horn astrocytes, while isolated running bouts did not. Future studies will need to determine the full extent (in rostrocaudal and dorsoventral directions), underlying mechanisms (e.g., descending noradrenergic, local glutamatergic signaling, gap junction involvement), behavioral dependencies (e.g., motor activity), and modulatory effects of this widespread astrocyte activity on spinal neural circuits and behavior (e.g., trans-segmental sensitization)^14-16^. Innocuous pinch evoked only sparse activity in isolated ROI clusters (**Fig. 2b**). The location of these clusters may provide clues about regional differences in spinal sensory processing and serves as a reminder that data from small FOVs can introduce sampling bias.

In summary, the technical capabilities, small form factor, and low weight of our wearable macroscopes pave the way for a wide range of exciting neuroscience applications. The repeatable and quantifiable stimuli employed in our study should facilitate comparative measurements in other transgenic mouse lines, such as recordings in genetically defined neuronal subpopulations to reveal their activity patterns in relation to astrocytes. Simultaneous measurement of multiple cell types’ activity patterns should be feasible given our macroscopes’ 450-570 nm achromatic range but will require custom multiband filters optimized for large incidence angles (**Methods**). By allowing measurements from large tissue areas and distributed cell networks in behaving animals, our macroscopes may find novel applications in various disease settings (e.g., structural/functional regeneration after spinal cord injury^17^) and larger experimental models (e.g., transgenic rats^7^, marmoset^18^). Our macroscope’s long working distance combined with implantable optics, such as microprisms (**Suppl. Fig. 3**), should enable high-speed translaminar measurement of sensorimotor activity from and across spinal laminae (e.g., related to motor control). Imaging during standard mouse behavioral assays should be feasible given the minimal mobility impediment mice showed during the open field test (**Suppl. Fig. 4**). Altogether, we expect end-users of our macroscopes to achieve real-time, high-resolution visualization of cellular activity patterns across large FOVs in a multitude of behavioral and disease contexts.

## Methods

### Miniature macroscope design

Our wearable macroscope includes six major components: the custom optics barrels (including eight microlenses), fluorescence filters, main body, image sensor module, illumination module, and base plate (**Fig. 1a**).

#### Optical design

The optical design was generated *de novo* using Zemax optical modeling software (Zemax LLC) and manual calculations. The final design form resembled a double Petzval-type short-conjugate relay. This fairly unique configuration allows for simultaneously large numerical aperture (NA) and field of view (FOV) while maintaining a reasonable principal ray angle (<15-degree half-cone incident on the dichroic and emission filters). Each half of the optical system included a negative element, fitting the Petzval-type description, which acts both as a field-flattener and to shorten the physical length of the optical system, akin to a telephoto configuration. During the design phase, geometric distortion was deliberately allowed to increase to 8.1% to gain improvement across the other Seidel aberrations (spherical, coma, astigmatism, and field curvature). The final optical design, which involved extensive global optimization, utilized eight lens elements made from five different glass materials (**Suppl. Table 1**) housed in two separate custom mechanical housings/barrels for easy assembly (Optics Technology Inc.). Both halves of the optical train were optimized together rather than independently, with each half compensating for the aberrations of the other to ensure high performance. To achieve near-uniform optical performance across the FOV, we incorporated a small field aperture at the plane of the primary emission filter, which allowed for an ∼37% improvement in resolution at the FOV edges while reducing light collection by only ∼11% (based on Zemax calculations). Light collection and resolution in the central FOV (∼2 mm diameter) were unaffected by this aperture. Based on Zemax calculation, the numerical aperture (NA) of our wearable macroscopes is 0.39.

Our macroscope uses a short-pass dichroic beamsplitter (FF496; Semrock Inc) and primary (ET525/50m; Chroma Technology Corp.) and secondary emission filters (FF01-496/LP; Semrock Inc.). These filters were chosen to maximize the emission signal from the green fluorescent genetically encoded calcium indicator GCaMP, currently the most advanced and common calcium reporter in use. No excitation filter was necessary because we employed a 473 nm DPSS laser as the excitation light source (MBL-FN-473; CNI Optoelectronics Tech. Co.).

Standard interference-type filters are designed for a particular angle of incidence (45 degrees for dichroic beamsplitters and 0 degrees for emission filters), with a tolerance of around ±5-7 degrees. To achieve a large FOV while maintaining reasonable device dimensions, we deliberately allowed a larger than usual principal ray incidence angle on the filters (±15 degrees). Increasing the incidence angle widens the passband of interference filters while shifting it toward shorter wavelengths. In particular, for the 45-degree dichroic beamsplitter, placed in the center of the optical train, light on the side closer to the image sensor reflects at a significantly larger angle than on the opposite side (60- and 30-degree incidence angle, respectively). This feature results in the dichroic beamsplitter’s spectral edge shifting by as much as ±25 nm around its nominal edge of λ=491 nm (at 45-degree incidence). On the side closer to the image sensor, the beamsplitter’s spectral edge shifts to λ=516 nm, rejecting more of the fluorescence signal, causing the corresponding image region to get darker. On the opposite side, the spectral edge shifts to λ=466 nm, accepting more stray excitation light, causing a brighter image region with lower contrast. For emission filters, the increased incidence angle results in the acceptance of more stray excitation light. We solved the stray light problem by placing a secondary emission filter in front of the image sensor, blocking most stray excitation light. The primary and secondary emission filters are approximately located at conjugate planes and, therefore, complement each other in angular performance.

The excitation light was delivered to the macroscope using a flexible multi-mode fiber (MMF) with a 200 μm core and 0.22 NA (FG200UEA; Thorlabs). To reduce laser speckle, we employed a square-core fiber (FP150QMT; Thorlabs) and a device that generates time-dependent mode-mixing within the MMF on a fast time scale through induced fiber bending (quick relative to the image acquisition speed). A square-core fiber was chosen because of its high degree of inherent mode-mixing and near isotropic mixing of modes over a large range of fiber-bend radii. A precision optical spacer and condenser lens (part no. 84-377; Edmund Optics) were used to collimate the illumination light. Note that optical fibers were chosen because they provide flexibility in coupling different light sources (e.g., DPSS lasers and LEDs), providing unique opportunities for future all-optical interrogations of cellular activity in behaving animals.

#### Optomechanical design

The macroscope housing was designed using CAD software (Autodesk Inventor) and included four modules (main body, sensor mount, illumination module, and base plate). The main body houses the imaging optics and fluorescence filters. The illumination module delivers the excitation light. The image sensor module mounts and provides heat dissipation for the CMOS sensor. The detachable base plate allows reproducible mounting, facilitating chronic imaging experiments. This modular design allows for modification of individual components without affecting the rest of the device (e.g., to upgrade the image sensor, incorporate an integrated LED instead of a fiber-coupled laser, or modify the baseplate to pursue an alternative mounting strategy).

The individual lens elements and primary emission filter were assembled inside two barrels to ensure light tightness. All fluorescence filters were edge-blackened (Sharpie Paint Pen). The dichroic cover, which also mounts the illumination module, was coated with highly absorptive black paint (Avian Black-S) to reduce reflections of back-scattered stray light.

Most of the housing was fabricated from light-weight black polyether ether ketone (PEEK), a thermoplastic opaque to visible and infrared (IR) light (near-IR light with an emission peak of around 850 nm was used to illuminate the behavioral setup during video recordings). The sensor mount was machined from aluminum to help dissipate heat generated by the image sensor, thereby reducing sensor noise. The housing was optimized over several iterations such that the fully assembled device weighed <10 g, ensuring that the animals could comfortably carry the macroscope on their back. We employed a folded imaging path to lower the device’s center of gravity and minimize torque on the animal (i.e., the short-pass dichroic beamsplitter reflected fluorescence emission and transmitted excitation light). Five-axis computer numerical control (CNC) milling was selected for ease of fabrication. This technique influenced the design process as the complex geometry needed to be confined to dimensions accessible to the CNC mill.

#### Electronic design

Fluorescence emission was detected with a CMOS sensor (MT9M024IX3STC, Aptina Imaging), which was chosen for its high-sensitivity (5.48 and 6.7 V/lux-sec for the RGB and monochrome versions, respectively), resolution (1280 × 960 pixels; 3.75 μm pixel spacing, corresponding to 2.63 μm at the object plane), and frame rate (45 fps at full resolution). Note that high frame rates are essential for spinal cord recordings given the rapid, large-amplitude, and non-uniform tissue displacements in this region during animal behavior^10^.

While the image sensor can record data at 14-bit (linear gamma), all data were recorded at 12-bit depth. The sensor was mounted on a custom miniature printed circuit board (PCB; Axelsys), interfaced with an Aptina data-acquisition board (AGB1N0CS-GEVK), and controlled using the Aptina DevWare Software Package.

#### Macroscope characterization

The macroscope’s optical performance (resolution and contrast) was determined by quantifying the device’s limit of resolution (LOR) (**Fig. 1**), point spread function (PSF) (**Suppl. Fig. 1**), and modulation transfer function (MTF) (**Suppl. Fig. 2**) before any software-based enhancements. The LOR (spatial frequency limit) was defined by the bar set where at least 5% contrast can be observed (the minimal contrast needed for detection by the human eye) and measured with the “Find Peaks” ImageJ plugin and custom ImageJ scripts. For PSF measurements, a point source was generated using a 1 μm pinhole (TC-RT01, Technologie Manufaktur), green fluorescence reference slide (2273, Ted Pella), and blue LED excitation source (M470L3, Thorlabs). The lateral and axial FWHM of the measured PSF was determined using Gaussian curve fits. The slight difference in our lateral LOR and PSF values is likely due to the practical limitations in measuring the PSF, which is the convolution of the optics’ impulse responses (note that the LOR test does not allow axial resolution measurements). To measure the modulation transfer function (MTF), we used the slanted edge test. Corresponding data analysis was performed using the “Slanted Edge MTF” ImageJ plugin, Excel, and GraphPad Prism. To quantify our device’s contrast limit, we chose the MTF10 metric (i.e., contrast at 10%). Although MTF is frequently reported at 0%, in our experience, MTF values below 10% are affected by sensor noise limits and optical phase reversal. Note that the slanted edge test allows for accurate measurements at spatial frequencies up to twice the Nyquist limit. Our MTF10 results with line widths less than the size of a single pixel (3 μm at the object plane) are well above measurement error. Line widths are equal to one-half the spatial period.

To characterize the macroscope’s FOV, we used a precision differential actuator (DRV3, Thorlabs) to image a grid target printed directly on the surface of a microscope slide (500 μm grid spacing) (**Fig. 1d**). While our current illumination module was designed to provide a flat illumination field, we observed moderate vignetting (i.e., image brightness reduction at the FOV edges) (**Fig. 1f**). Apart from the vignetting common to large FOV imaging systems, this effect resulted from a field stop we deliberately introduced for improving image quality at the FOV edges. To compensate for this synthetic vignetting and enhance the signal-to-noise ratio (SNR), an illumination module with a non-flat illumination field will need to be designed. This non-trivial engineering task may involve using multiple optical elements, non-imaging type optics, or miniature LEDs directly attached to the underside of the macroscope or baseplate. Currently, we correct for vignetting with post-acquisition software. While this introduces digital noise (due to the effectively reduced sensor bit depth at the FOV edges), this noise was not prominent in our *in vivo* recordings because of the large ΔF/F signals we typically observed in live animals (**Fig. 2d-f**).

The macroscope’s working distance was measured by placing the device in contact with a grid target (R1L3S3P, Thorlabs) and then translating it upwards until the image came into focus, using a precision differential actuator (DRV3, Thorlabs). Note that the working distance depends on the cover glass thickness. We used a #0 cover glass (∼100 μm thick) for our experiments.

Tissue analogs were used to estimate attainable imaging depth (**Suppl. Fig. 3**). These tissue phantoms consisted of 0.5% (by weight) agarose, 11% (by volume) scattering bead solution (1.00 μm Polybead Microspheres; part no. 07310-15; Polysciences), and 5% (by volume) fluorescent bead solution (6 μm FocalCheck Microspheres; part no. F14807; ThermoFisher), prepared using deionized water. Contrast and FWHM in **Suppl. Fig. 3g-j** were measured without background subtraction using the ImageJ plugin MetroloJ.

The integrated device weight was determined by weighing the individual components and the assembled system. We did not include the sensor cabling wires or optical fiber in the measurement as the weight of these is typically supported by external mounts or a commutator.

### Experimental model and subject details

All procedures were performed following the National Institutes of Health (NIH) guidelines and were approved by the Institutional Animal Care and Use Committee (IACUC) at the Salk Institute. Mouse strains used in this study included GFAP-Cre (RRID: IMSR_JAX:012886), Ai95(RCL-GCaMP6f)-D (RRID: IMSR_JAX:024105), and Cx3cr1^+/eGFP^ mice (RRID: IMSR_JAX:005582). Mice were group-housed, provided with bedding and nesting material, and maintained on a 12-h light-dark cycle in a temperature (22 ± 1°C) and humidity controlled (45-65%) environment. All the imaging and behavioral experiments involved 7-9 weeks-old heterozygous female mice. Experimental mice used in individual experiments typically originated from different litters. Mice had marks for unique identification. No criteria were applied to allocate mice to experimental groups.

### Live animal preparation

Animals were implanted with a spinal and head plate under general anesthesia approximately one week before laminectomy, as previously described^10^. Buprenex SR (0.5 mg/kg) was given to minimize post-operative pain.

A laminectomy (typically 2 wide × 4 mm long) was performed at the T12-T13 vertebra level, corresponding to spinal segments L2-L5^4^, four to seven days before the initial recording session. The dura mater overlying the spinal cord was kept intact, and a custom-cut #0 coverslip was used to seal the laminectomy creating an optical window for imaging. The coverslip was replaced immediately before recording sessions to maximize optical clarity.

### Fluorescence imaging

#### *In vivo* imaging through a dorsal optical window

Data in behaving mice were acquired following previously established protocols^10^. A fiber-coupled 473 nm DPSS laser was used for GCaMP6f excitation. Approximately 10-20 recordings were taken per imaging session, with each recording lasting around 1-2 min. Recordings were acquired at ∼25-75 μm focal depth below the pia. The typical average power for a given recording was <105 μW mm^-2^. No signs of phototoxicity, such as a gradual increase in baseline fluorescence, lasting changes in activity rate, or blebbing of labeled cells, were apparent in our recordings. All *in vivo* data were acquired at the image sensor’s full resolution (1,280 × 960 pixels) and maximum frame rate (45 Hz).

#### *In vitro* imaging through implanted microprisms

To demonstrate the wearable macroscopes’ long working distance, we implanted 2.0 mm × 2.0 mm × 2.0 mm microprisms (W × D × H) in tissue phantoms with embedded 6 μm-diameter fluorescent beads (**Suppl. Fig. 3**). The miniature macroscope’s FOV allowed imaging of the entire microprism face. Recordings were taken at different focal depths from the vertical microprism-tissue phantom interface, where 0 µm was defined as the point when fluorescent beads first came into focus. A fiber-coupled 473 nm DPSS laser was used for fluorescence excitation.

#### Widefield imaging of tissue sections

To benchmark our wearable macroscope’s imaging performance, we used a commercial benchtop macroscope with similar FOV (**Fig. 1e-f**). This macroscope was equipped with a wide-FOV objective (XL Fluor4x/340; Olympus Corp.) and a high-resolution image sensor (ORCA FLASH4.0 V3, Hamamatsu Photonics K.K.). Despite being orders of magnitude smaller and lighter, our wearable macroscopes provided comparable resolution and contrast and ∼2x better light collection due to their higher NA of 0.39 compared to the commercial system’s 0.28.

### Sensory stimuli and behavioral tests

#### Sensory stimuli during *in vivo* imaging

Each imaging session consisted of up to 20 recordings. Mechanical stimuli were delivered to the animal’s tail using a rodent pincher system (cat. no. 2450; IITC Life Science, Inc.). Pinch pressures were applied in the dorsoventral direction at approximately 6 mm from the base of the animal’s tail. Each pinch stimulus (typically one per recording) lasted around 1-2 s. Subsequent stimuli were delivered at least 1.5-2 minutes apart to minimize response adaptation. The order in which stimuli of different amplitudes were delivered was randomized. **Fig. 2** shows data from p < 200 g and p > 500 g mechanical stimuli.

#### Open-field test

To quantify how the ∼9.8 g macroscope mounted on the animal’s back (lumbar spinal cord) might affect general locomotor activity, we used the open-field test with infrared tracking (ENV-510; Med Associates Inc.) (**Suppl. Fig. 4**). Animals (∼8-weeks-old female GFAP-GCaMP6f mice with an implanted spinal plate) were acclimated to the testing environment by placing their home cage in the testing room ∼60 minutes before the recordings. Mice were then transferred to individual testing chambers, always placed in the lower-left corner facing the center of the apparatus. Six five-minute recordings were collected for each animal (N=3) with the experimenter outside the room. The mice were then removed from the apparatus, the wearable macroscope mounted, and placed back in their respective chambers using the same starting orientation. Another six five-minute recordings were acquired with the experimenter outside the room. Average running speed (cm/s), total distance traveled (cm), and rearing activity (vertical movement count) were analyzed. Recordings of a given animal were averaged before averaging across animals.

### Image data processing and analysis

#### *In vivo* imaging data

We used a custom Matlab script to convert the image sensor’s raw files from 12-to 16-bit and adjust their pixel range. The data were then further processed in Fiji. We converted the image data to TIFF, cropped the time-lapse recordings to the laser-on period and fluorescently labeled central areas of the FOV, followed by computational illumination correction and background subtraction. Full-frame image motion was reduced using Moco^19^. Image edge artifacts introduced by motion correction were cropped. Within-frame distortions were typically small at the employed sampling rate (45 Hz) and therefore not corrected. Recordings for which motion correction failed (e.g., vigorous running of >100 mm/s) were excluded from data analysis.

Motion-corrected calcium imaging data were analyzed using custom ImageJ and MATLAB software. We used an unbiased tiling approach to quantify calcium activity across the FOV, similar to our previous work^20^ (**Fig. 2**). Each tile corresponded to a 4 × 4-pixel (equivalent to ∼10 × 10 μm) ROI. We defined three ROI classes: over, near, and distant from blood vessels. First, we calculated the mean and s.d. of the recording’s maximum intensity image. Next, we determined individual ROIs’ average intensity values (A). If A < mean - 1 s.d., it was considered a blood vessel-ROI. If A ≥ mean + (1+x)*s.d. (with × ∈ [0-1.2] depending on data set (e.g., blood vessel size/pattern)), it was classified as distant from blood vessels. ROIs with in-between values were considered near-blood vessel-ROIs. ROIs over and near blood vessels were excluded from data analysis. For all ROIs distant from blood vessels, we calculated their average fluorescence intensity over time. Corresponding activity traces were temporally smoothed using an 0.4 s sliding average. Matlab’s “findpeaks” function identified local maxima within the traces. The fluorescence baseline and noise level (in s.d.) of a given trace were calculated using a 2 s period before stimulus onset. Local maxima were considered evoked activity if fluorescence intensity values surrounding the peak were at least 6 s.d. above baseline for ≥2 s. Calcium transient onset was defined as the point at which the fluorescence intensity trace immediately before the peak crossed a 2 s.d. threshold above baseline. Calcium transient duration was defined as the trace’s full width at half maximum (FWHM). Traces that showed a) activity within 2 s after recording onset (i.e., before stimulus onset), b) fluorescence decreases faster than the indicator’s unbinding kinetics (≥50% signal drop within ≤250 ms), or c) a 6 s.d. drop below fluorescence baseline after transient onset for ≥3 s were considered artifactual (e.g., caused by tissue motion) and excluded from further analysis. For multi-peak transients, the largest peak’s value was used for FWHM calculation. If the intensity values between peaks fell below the half-maximum value for ≥250 ms, the half-maximum point closest to the largest peak was used for FWHM calculation. If the trace’s amplitude fell below 2 s.d. above baseline for ≥400 ms, the corresponding transients were considered separate calcium spikes.

To distinguish spontaneous from evoked calcium activity, we applied additional criteria. A calcium transient was considered evoked if its onset occurred within 5 s after stimulus onset. For all activity traces that passed the filters mentioned above, we calculated ΔF(t)/F_0_(t). F_0_ was determined with Matlab’s mode function using a 15-bin width. Traces that showed abrupt intensity changes (≥0.5 ΔF/F within ≤65 ms) were removed from the analysis.

To quantify potential differences in sensory-evoked activity on the left and right sides of the spinal cord (**Fig. 2j, k**), we focused our analysis on 100 μm-wide gray matter regions (±150-250 μm distance from the central vein).

### Analog and video data processing and analysis

All analog data were synchronously recorded at 1 kHz using DAQExpress 2.0 software (National Instruments). Analog data included the pressure sensor output from the rodent pincher system, treadmill speed, and on-off TTL signal of the miniaturized macroscope’s light source, depending on the experiment. Pinch application and mouse behavior were also recorded on a video camera (≥20 Hz; Stingray F-033, Allied Vision Technologies). To synchronize imaging with video data, we placed a near-infrared LED within the video camera’s FOV, triggered from the wearable macroscope’s light source drive signal (TTL pulse). Imaging and analog data were synchronized by recording the on-off TTL signal of the macroscope’s light source together with all other analog data. To relate the animal’s locomotor activity more closely to the imaging data, we focally restrained animals on a spherical treadmill equipped with an optical encoder (E7PD-720-118, US Digital), allowing precise readout of running speed. All analog data was processed using custom MATLAB (Mathworks) routines. The pincher and encoder traces were first cropped to the light source-on period. Pressure traces were then quantified with respect to stimulus amplitude and duration. Encoder traces were smoothed using a sliding average (window size: 0.4 s). Locomotion onset or offset was defined as the point at which the smoothed running speed exceeded or fell below 10 mm/s. Encoder traces were analyzed concerning running speed, duration, and frequency. If the running speed fell below the 10 mm/s threshold for ≥750 ms, the local maxima were considered separate running bouts. The video data were cropped to the LED-on period. Videos were scored manually regarding pinch onset and offset. Calcium transient latency was calculated based on these measurements because video recordings provided higher temporal resolution than the pincher traces. For population analysis, data was computationally sorted using custom MATLAB (Mathworks) routines. Only trials with 1.5 ± 0.5 s pinch duration were included in the analysis.

### Statistical analysis

All data were analyzed and plotted using MATLAB, Excel, or GraphPad Prism software. Paired t-tests were used to evaluate potential calcium activity differences on the left and right sides of the spinal cord (**Fig. 2j, k**) and changes in the animal’s general locomotor activity (**Suppl. Fig. 4**). All data are represented as mean ± s.e.m. Group sample sizes were chosen based on power analysis or previous studies. The following convention was used to indicate *P* values: ‘NS’ indicates *P*>0.05, ‘*’ indicates 0.01<*P*≤0.05, ‘**’ indicates 0.001<*P*≤0.01, ‘***’ indicates 0.0001<*P*≤0.001, and ‘****’ indicates *P*≤0.0001.

### Reporting summary

Further information on research design is available in the Research Reporting Summary linked to this paper.

### Data availability

The data that support the findings of this study will be deposited in the Brain Image Library (BIL; https://www.brainimagelibrary.org/index.html). They will also be available from the corresponding author upon reasonable request.

### Code availability

The custom Java- and Matlab-based code used to process and analyze the data will be deposited in GitHub. It will also be available from the corresponding author upon reasonable request.

## Acknowledgments

We thank members of the Nimmerjahn lab for comments on the manuscript, the Salk machine shop and Nick Andrews of the Behavior Testing Core for technical support, and J. Chambers for mouse colony management. This work was primarily supported by the National Institutes of Health (NIH) grant R01NS108034 (A.N). It was partially supported by the NIH grants U01NS103522, U19NS112959, U19NS123719, a Salk Innovation Grant, and equipment funds from C. and L. Greenfield (A.N.). P.S. was supported by a Rose Hills Foundation graduate fellowship and N.A.N. by funds from an NIH T32/CMG Training Grant, Burt and Ethel Aginsky Research Scholar Award, Kavli-Helinski Endowment Graduate Fellowship, and NIH individual predoctoral fellowship (F31NS120619). The content is solely the authors’ responsibility and does not necessarily represent the official views of the NIH.

## Author contributions

P.S., D.D., and A.N. conceived and designed the study with input from E.M.C. and N.A.N. P.S. developed and characterized the wearable macroscopes and wrote ImageJ-based data analysis code. D.D. performed the *in vivo* imaging experiments. E.M.C. conducted the motor behavior experiments. P.S., D.D., and E.M.C. analyzed the *in vitro* and *in vivo* data. A. Ngo, G.G., and J.A.O. developed Matlab- and ImageJ-based data analysis code. A.N. supervised the study and wrote the initial manuscript draft. All authors contributed to the text and figures, discussed the results, or provided input and edits on the manuscript.

## Competing interests

The authors declare no competing interests.

Correspondence and requests for materials should be addressed to A.N.

## Supplementary Figures

**Suppl. Fig. 1.**
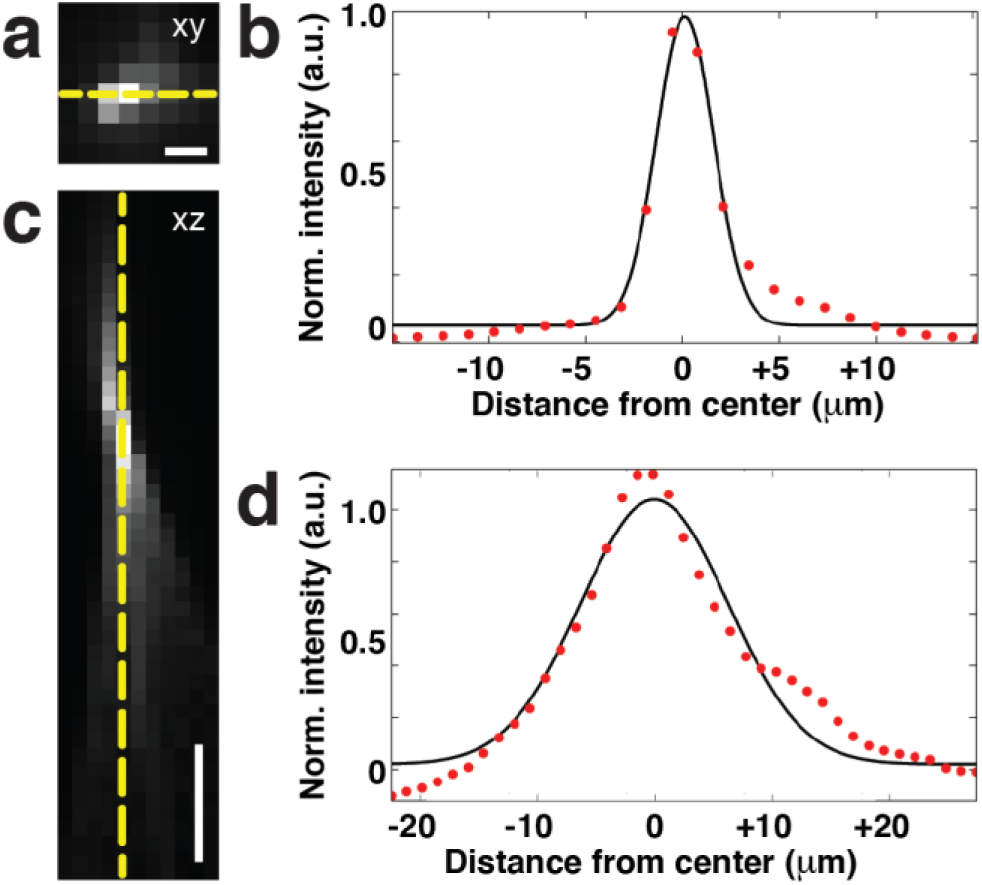
Wearable macroscopes with custom-compound microlenses provide ∼3 μm lateral resolution. **a**, Experimentally measured lateral point spread function (PSF) of the integrated wearable macroscope. Scale bar, 5 μm. **b**, × cross-section showing a full width at half maximum (FWHM) of ∼3.3 μm. **c**, Experimentally measured axial PSF of the integrated wearable macroscope. Scale bar, 20 μm. **b**, z cross-section showing an FWHM of ∼17.7 μm. The red dotted and black solid lines in b and d show the measured intensity (along the yellow dashed lines in a and c) and Gaussian fit profile, respectively.

**Suppl. Fig. 2.**
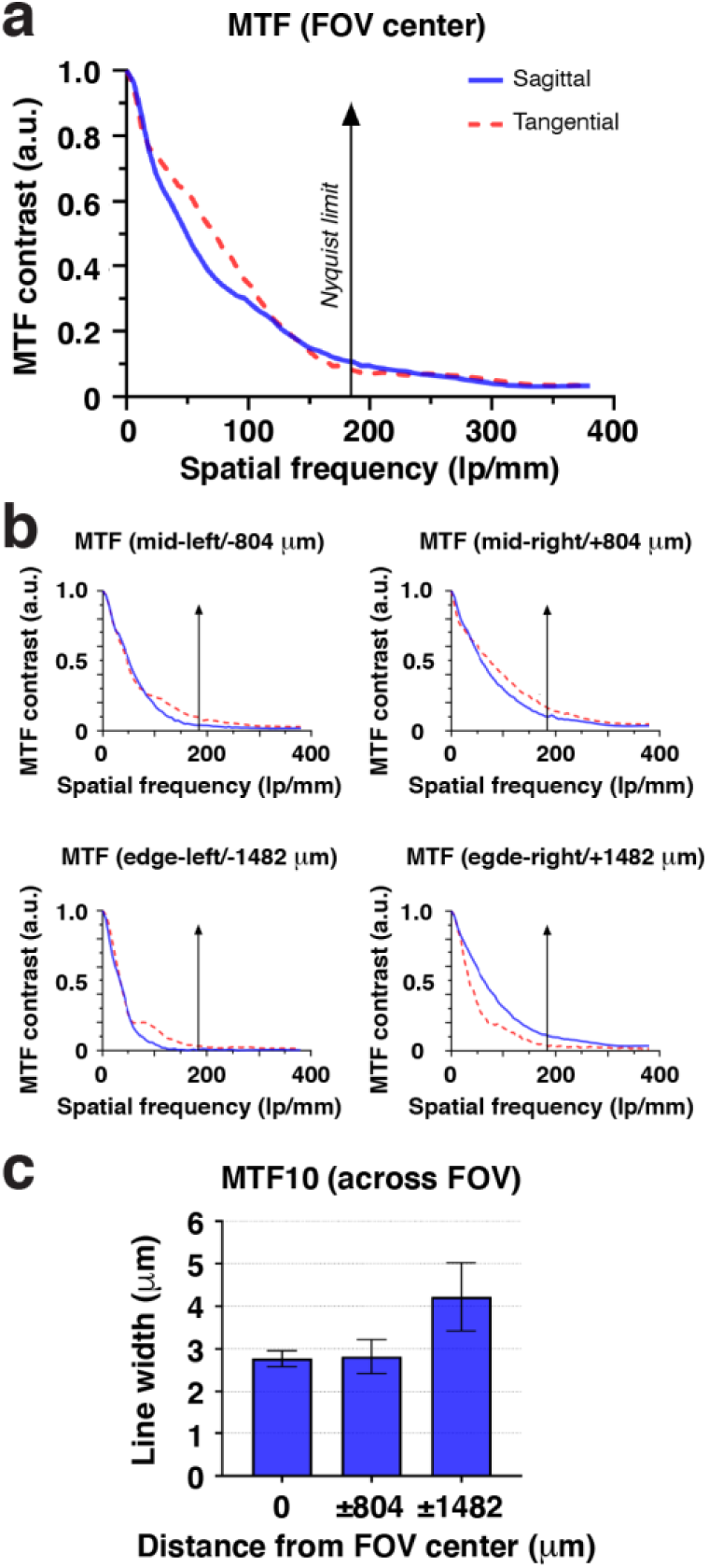
Wearable macroscopes with custom-compound microlenses provide high contrast across the field of view. **a**, Modulation transfer function (MTF) of the integrated macroscope measured in the center of the field of view (FOV) using the Slanted Edge test. **b**, MTF at different indicated FOV positions relative to the center. **c**, MTF contrast at 10% (MTF10) across the FOV. Displayed values are averages across similar FOV locations and horizontal and vertical Slanted Edge targets. Spatial frequencies were converted to line widths. The data are presented as mean ± s.e.m. The larger error bars toward the FOV edge likely indicate sample tilt.

**Suppl. Fig. 3.**
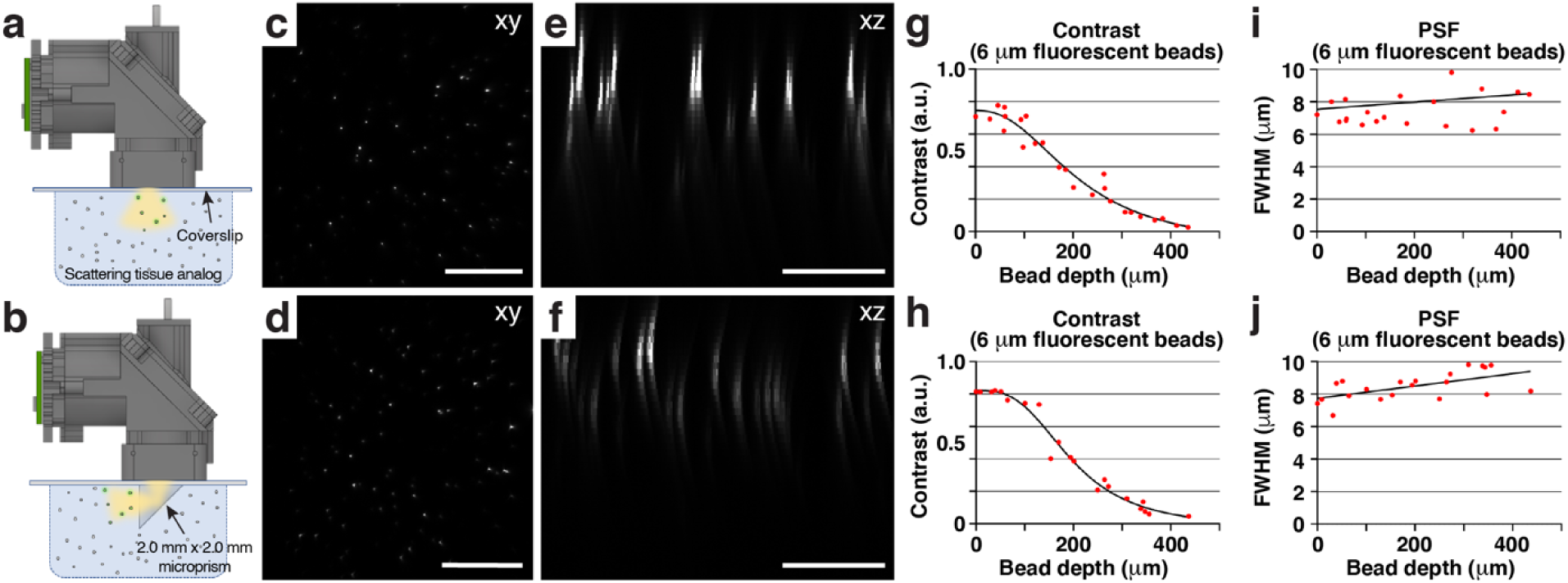
Wearable macroscopes with custom-compound microlenses permit imaging through implanted microprisms. **a-b**, Schematics showing the experimental approach for characterizing and comparing two imaging conditions in scattering tissue phantoms (**Methods**): *Top*, imaging through a coverslip; *bottom*, imaging through a coverslip with an attached 2.0 mm × 2.0 mm × 2.0 mm microprism (W × D × H). **c-d**, Example images of tissue phantom embedded 6 μm-diameter fluorescent beads. Each image is a maximum intensity projection through a z-stack acquired as shown in a-b by translating the wearable macroscope axially. Scale bar, 500 μm. **e-f**, Maximum intensity side projections of the acquired z-stacks. Scale bar, 200 μm. **g-h**, Bead contrast as a function of imaging depth. **i-j**, Lateral full width at half maximum (FWHM) of the 6 μm-diameter fluorescent beads as a function of imaging depth.

**Suppl. Fig. 4.**
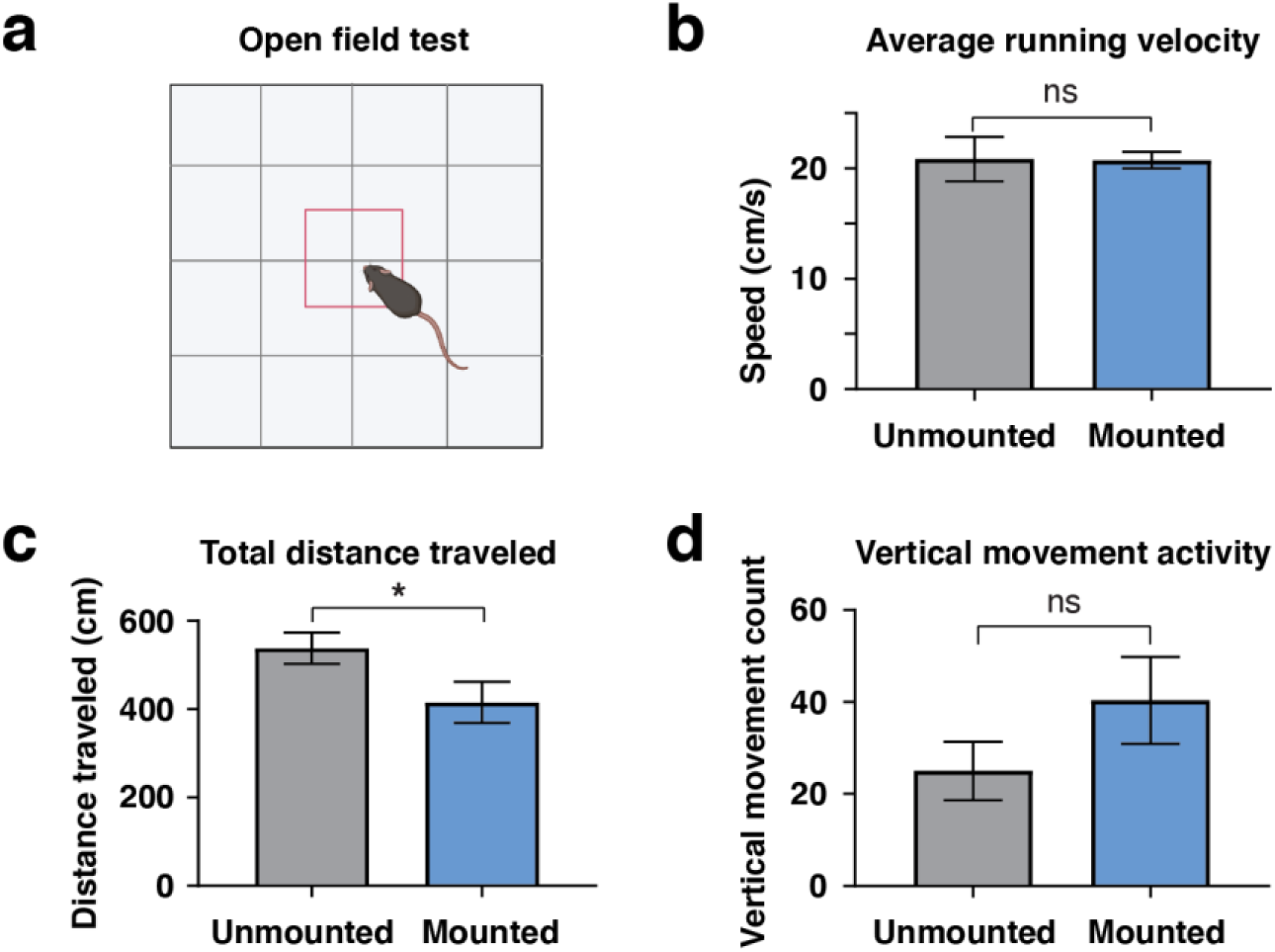
Spine-mounted macroscopes of <10 grams have little effect on mouse open-field behavior. **a**, Schematic of the open-field test used to compare the animal’s general locomotor activity with or without the ∼9.8 g macroscope mounted over its lumbar spinal cord. **b-d**, Population data showing the average running speed (b), total distance traveled (c), and vertical movement/rearing activity (d) before and after wearable macroscope mounting. The data in b-d are from six five-minute recordings before and after attaching the device. All animals (N=3) were tested before and after macroscope mounting (**Methods**). Paired t-tests determined *P* values, and all data are presented as mean ± s.e.m.

## Supplementary Tables

**Suppl. Table 1.** Wearable macroscope optical design.

| Surface | Type | Radius | Thickness | Glass | Diameter |
| --- | --- | --- | --- | --- | --- |
| OBJ | STANDARD | Infinity | 0.40000 | N-BK7 | 6.00000 |
| 1 | STANDARD | Infinity | 3.83988 |  | 5.95166 |
| 2 | STANDARD | -5.74334 | 1.60002 | N-SF66 | 5.36800 |
| 3 | STANDARD | 15.70964 | 0.81873 |  | 6.41800 |
| 4 | STANDARD | -32.74580 | 1.60104 | S-LAH97 | 6.89400 |
| 5 | STANDARD | -8.05357 | 0.21700 |  | 7.61200 |
| 6 | STANDARD | Infinity | 1.60035 | N-LAF34 | 8.46000 |
| 7 | STANDARD | -12.51519 | 0.52205 |  | 8.77400 |
| 8 | STANDARD | 21.51710 | 1.60150 | S-LAH55V | 9.00000 |
| 9 | STANDARD | -47.46657 | 0.50000 |  | 9.00000 |
| 10 | STANDARD | Infinity | 1.00000 | FSILICA | 8.92000 |
| 11 | STANDARD | Infinity | 5.50000 |  | 8.81600 |
| 12 | COORDBRK | - | 0.00000 |  | - |
| 13 | STANDARD | Infinity | 0.00000 | MIRROR | 12.21416 |
| 14 | COORDBRK | - | 5.50000 |  | - |
| STO | STANDARD | -33.34728 | 1.98940 | S-LAH97 | 7.12400 |
| 16 | STANDARD | 6.13755 | 1.00000 | N-SF66 | 7.23800 |
| 17 | STANDARD | 17.95736 | 0.10000 |  | 7.75400 |
| 18 | STANDARD | -6.35500 | 1.89487 | S-LAH58 | 8.06200 |
| 19 | STANDARD | -35.88850 | 1.60925 |  | 7.63800 |
| 20 | STANDARD | Infinity | 0.74100 |  | 6.07952 |
| 21 | STANDARD | 15.94777 | 0.99962 | N-SF66 | 5.53400 |
| 22 | STANDARD | -21.55705 | 2.80317 |  | 5.09800 |
| 23 | STANDARD | Infinity | 0.10000 | N-BK7 | 4.08338 |
| IMA | STANDARD | Infinity |  |  | 4.05844 |

## Supplementary Movies

**Suppl. Movie 1** | **High-speed trans-segmental imaging of innocuous tail pinch-evoked calcium activity in behaving GFAP-GCaMP6f mice**. Example time-lapse recording acquired with the wearable macroscope showing innocuous tail pinch-evoked calcium excitation in spinal astrocytes. The data were obtained at ∼45 fps and ∼50 μm focal depth below the pia. Elapsed time is indicated in the upper right corner (total duration: 35 s). To allow the precise readout of locomotor activity, the mouse was placed on a spherical treadmill. Innocuous pinch triggered sparse calcium excitation across the imaged lumbar spinal segments. Running alone did not evoke significant calcium increases. Scale bar, 250 μm.

**Suppl. Movie 2** | **High-speed trans-segmental imaging of noxious tail pinch-evoked calcium activity in behaving GFAP-GCaMP6f mice**. Example time-lapse recording acquired with the wearable macroscope showing noxious tail pinch-evoked calcium excitation in spinal astrocytes. The data were obtained at ∼45 fps and ∼50 μm focal depth below the pia. Elapsed time is indicated in the upper right corner (total duration: 34 s). To allow the precise readout of locomotor activity, the mouse was placed on a spherical treadmill. Noxious pinch triggered widespread, coordinated calcium excitation on both sides of the spinal cord and across lumbar spinal segments. Scale bar, 250 μm.

## References

1 Koch, S. C., Acton, D. & Goulding, M. Spinal Circuits for Touch, Pain, and Itch. Annu Rev Physiol 80, 189–217, doi:10.1146/annurev-physiol-022516-034303 (2018).

2 Nelson, N. A., Wang, X., Cook, D., Carey, E. M. & Nimmerjahn, A. Imaging spinal cord activity in behaving animals. Exp Neurol 320, 112974, doi:10.1016/j.expneurol.2019.112974 (2019).

3 Abraira, V. E. & Ginty, D. D. The sensory neurons of touch. Neuron 79, 618–639, doi:10.1016/j.neuron.2013.07.051 (2013).

4 Harrison, M. et al. Vertebral landmarks for the identification of spinal cord segments in the mouse. Neuroimage 68, 22–29, doi:10.1016/j.neuroimage.2012.11.048 (2013).

5 Aharoni, D. & Hoogland, T. M. Circuit investigations with open-source miniaturized microscopes: Past, present and future. Front Cell Neurosci 13, 141, doi:10.3389/fncel.2019.00141 (2019).

6 Chen, S. et al. Miniature fluorescence microscopy for imaging brain activity in freely-behaving animals. Neurosci Bull 36, 1182–1190, doi:10.1007/s12264-020-00561-z (2020).

7 Scott, B. B. et al. Imaging Cortical Dynamics in GCaMP Transgenic Rats with a Head-Mounted Widefield Macroscope. Neuron 100, 1045–1058 e1045, doi:10.1016/j.neuron.2018.09.050 (2018).

8 Rynes, M. L. et al. Miniaturized head-mounted microscope for whole-cortex mesoscale imaging in freely behaving mice. Nat Methods 18, 417–425, doi:10.1038/s41592-021-01104-8 (2021).

9 Guo, C. et al. Miniscope-LFOV: A large field of view, single cell resolution, miniature microscope for wired and wire-free imaging of neural dynamics in freely behaving animals. bioRxiv, doi: https://doi.org/10.1101/2021.1111.1121.469394 (2021).

10 Sekiguchi, K. J. et al. Imaging large-scale cellular activity in spinal cord of freely behaving mice. Nat Commun 7, 11450, doi:10.1038/ncomms11450 (2016).

11 Kohro, Y. et al. Spinal astrocytes in superficial laminae gate brainstem descending control of mechanosensory hypersensitivity. Nat Neurosci 23, 1376–1387, doi:10.1038/s41593-020-00713-4 (2020).

12 Xu, Q. et al. Astrocytes contribute to pain gating in the spinal cord. Sci Adv 7, eabi6287, doi:10.1126/sciadv.abi6287 (2021).

13 Nam, Y. et al. Reversible Induction of Pain Hypersensitivity following Optogenetic Stimulation of Spinal Astrocytes. Cell Rep 17, 3049–3061, doi:10.1016/j.celrep.2016.11.043 (2016).

14 Merten, K., Folk, R. W., Duarte, D. & Nimmerjahn, A. Astrocytes encode complex behaviorally relevant information. bioRxiv, doi: https://doi.org/10.1101/2021.1110.1109.463784 (2021).

15 Orts-Del’Immagine, A., Dhanasekar, M., Lejeune, F. X., Roussel, J. & Wyart, C. A norepinephrine-dependent glial calcium wave travels in the spinal cord upon acoustovestibular stimuli. Glia, doi:10.1002/glia.24118 (2021).

16 Mu, Y. et al. Glia Accumulate Evidence that Actions Are Futile and Suppress Unsuccessful Behavior. Cell 178, 27–43 e19, doi:10.1016/j.cell.2019.05.050 (2019).

17 Ceto, S., Sekiguchi, K. J., Takashima, Y., Nimmerjahn, A. & Tuszynski, M. H. Neural Stem Cell Grafts Form Extensive Synaptic Networks that Integrate with Host Circuits after Spinal Cord Injury. Cell Stem Cell 27, 430–440 e435, doi:10.1016/j.stem.2020.07.007 (2020).

18 Kondo, T. et al. Calcium Transient Dynamics of Neural Ensembles in the Primary Motor Cortex of Naturally Behaving Monkeys. Cell Rep 24, 2191–2195 e2194, doi:10.1016/j.celrep.2018.07.057 (2018).

## References

19 Dubbs, A., Guevara, J. & Yuste, R. moco: Fast Motion Correction for Calcium Imaging. Front Neuroinform 10, 6, doi:10.3389/fninf.2016.00006 (2016).

20 Patriarchi, T. et al. Ultrafast neuronal imaging of dopamine dynamics with designed genetically encoded sensors. Science 360, doi:10.1126/science.aat4422 (2018).

